# AffectRoute: Role-Structured EEG and Peripheral Physiology for Subject-Independent Affect Regression

**DOI:** 10.64898/2026.08.24.746535

**Authors:** Ruiyang Zhang, Xianglian Jia

## Abstract

Subject-independent affect regression from physiological signals remains challenging because emotional responses vary substantially across individuals, widely used datasets provide only coarse trial-level annotations, and heterogeneous physiological modalities may not contribute reliably when treated as if they were interchangeable predictors. We have developed AffectRoute, a protocol-conditioned subject-independent affect regression that is conditioned on the protocol and assigns separate predictive functions to information from the population, electroencephalography (EEG), and peripheral physiological signals (PPS). First, a source-population prior establishes a trial-level affective anchor using only the data from source participants. TrajBridge then combines an EEG representation that is supervised by REFED for participant-specific adjustments with temporal structure obtained from the continuous REFED annotations in order to create a weakly supervised segment-resolved pseudo-trajectory and to establish a frozen trial-level baseline. PhysioRoute next reduces the remaining error by breaking down the residual correction into a source-derived direction, which is estimated from the out-of-fold residuals within the source group, and a channel-specific magnitude derived from the PPS. When evaluated on DEAP and DREAMER using a leave-one-subject-out approach at the participant level, AffectRoute showed consistent step-by-step improvements in both the mean absolute error and the concordance correlation coefficient. A method that relied solely on the source data was clearly worse than PhysioRoute, showing that the final improvement cannot be accounted for by transferable source residual regularity alone. Conventional alternatives to fusing the PPS were also found to be consistently less effective, although analyses at the channel level and with a leave-one-channel-out design showed that the peripheral contributions are axis-dependent yet distributed across channels. These results indicate that structured residual inference is an effective alternative to unrestricted multimodal fusion for subject-independent affect regression.

## 1. Introduction

Emotion recognition from physiological signals is an important component of affective computing, brain-computer interfaces, and human-centered intelligent systems because physiological responses can reveal affective information that is not always apparent from observable behavior. Electroencephalography (EEG) is particularly attractive because it captures neural activity with high temporal resolution, while peripheral physiological signals (PPS), including electrodermal, electromyographic, respiratory, cardiovascular, eye-movement, and temperature signals, provide complementary information about autonomic and somatic responses. In subject-independent settings, affect models are required to predict participants whose affective ratings are unavailable during training. Subject-independent prediction is therefore particularly challenging because physiological responses vary across individuals in baseline level, response magnitude, and distribution [1]. This issue is especially relevant to widely used datasets such as DEAP [2] and DREAMER [3], where participant-wise evaluation provides a natural setting for assessing performance on unseen participants.

A second challenge arises from the coarse supervision available in DEAP and DREAMER. Both datasets provide retrospective valence and arousal ratings at the trial level but do not provide continuous or segment-level affect annotations within each trial. Assigning the same trial rating to every EEG segment removes possible within-trial variation, whereas estimating an unconstrained segment-level sequence from a single post-stimulus label is inherently ambiguous. Multiple-instance and weakly supervised methods have addressed related coarse-to-fine affect-recognition settings, showing that trial-level supervision can be exploited to learn finer representations without requiring dense labels [4,5]. Furthermore, the REFED dataset [6] provides continuous real-time valence and arousal annotations together with synchronized EEG, offering an external source of fine-grained supervision for learning affect-related EEG representations and temporal organization. This external supervision can be transferred while DEAP and DREAMER remain trained and evaluated under their original trial-level annotation setting. Recent naturalistic-viewing studies further suggest that affective dynamics, particularly arousal-related dynamics, can generalize across participants and films, supporting the use of externally learned temporal structure as a meaningful constraint rather than an arbitrary synthetic fluctuation [7].

A third challenge concerns how EEG and PPS should be combined. Conventional multimodal emotion-recognition methods typically combine physiological modalities through feature-, representation-, or decision-level fusion, while recent architectures use modality-specific encoders and attention-based fusion [8,9]. These studies demonstrate the complementarity of EEG and PPS, but they also require a joint predictor to determine how heterogeneous modalities should contribute to the complete affect estimate. This requirement is non-trivial because EEG and peripheral channels arise from different physiological mechanisms and temporal scales, while the reliability of peripheral measurements can vary with sensor placement, motion, and acquisition conditions [10-12]. Moreover, peripheral channels are not uniformly informative across affective dimensions: facial electromyographic activity, electrodermal responses, respiration, cardiovascular signals, and temperature capture different aspects of emotional responding. These factors motivate a role-structured alternative in which EEG and PPS are not forced to act as interchangeable full predictors.

To address these three challenges, we propose AffectRoute, a protocol-conditioned, subject-independent affect regression framework comprising three sequential stages: (1) a source-population trial prior, (2) TrajBridge, and (3) PhysioRoute. A source-population trial prior first provides an affective anchor for each prespecified trial coordinate using source participants only. TrajBridge then constructs an EEG-centered pseudo-trajectory through two complementary forms of REFED transfer: a REFED-supervised EEG representation supports participant-specific center adjustment, while a temporal component derived directly from continuous REFED annotations provides externally supervised within-trial organization. The TrajBridge estimate therefore combines the source-population trial center with the EEG-based participant adjustment and the REFED-derived temporal component. PhysioRoute subsequently models the prediction error that remains after TrajBridge. For source participants, this residual is the difference between the observed trial rating and the corresponding TrajBridge prediction. Source-inner out-of-fold residuals at the same trial coordinate determine whether the TrajBridge estimate tends to require an upward or downward correction across participants, while source-trained channel-specific PPS models estimate candidate magnitudes for this remaining residual rather than predicting the complete affective target. For an unseen participant, the final AffectRoute prediction is obtained by adding the PPS-derived correction magnitude, with its sign determined by the source-derived residual direction, to that participant’s TrajBridge prediction. This sequential construction assigns each information source a distinct predictive role rather than combining EEG and PPS through unrestricted representation fusion.

### The main contributions of this work are threefold

1. A protocol-conditioned subject-independent formulation. AffectRoute performs participant-wise leave-one-subject-out prediction while using only source-participant ratings to construct the trial prior. The held-out participant contributes observable EEG, available PPS, and the prespecified trial coordinate, while that participant’s affect ratings remain unavailable until final evaluation.
2. TrajBridge for externally supervised pseudo-trajectory construction. TrajBridge separates estimation of the trial-level affective center from specification of within-trial temporal variation. REFED contributes both an EEG representation for participant-specific center adjustment and an annotation-derived temporal component learned from continuous affect supervision from REFED, allowing coarse target-dataset ratings to be expanded into structured pseudo-trajectories without imposing uniform segment labels.
3. PhysioRoute for channel-specific peripheral residual correction. Rather than using PPS as a second complete affect predictor, PhysioRoute models the residual that remains after TrajBridge. Same-trial source-inner OOF residuals determine whether the TrajBridge prediction should be corrected upward or downward, while source-trained channel-specific PPS models estimate the magnitude of that correction. The resulting signed residual correction is added to the TrajBridge-prediction. This direction-magnitude decomposition separates transferable cross-subject correction structure from peripheral physiological evidence and avoids both a fixed sensor hierarchy and unrestricted multimodal fusion.

We evaluate AffectRoute on DEAP and DREAMER under subject-independent leave-one-subject-out protocols. The experiments examine stage-wise gains from the population prior, TrajBridge, and PhysioRoute; compare constrained routing with conventional residual-fusion alternatives; assess channel-level PPS contribution through leave-one-channel-out analysis.

## 2. Methods

### 2.1 Datasets and Subject-Independent Protocol

DEAP and DREAMER were used as the target datasets for subject-independent affect regression, whereas REFED served as an external source of fine-grained supervision for EEG representation learning and temporal organization. [2,3,6] DEAP contains EEG and peripheral physiological recordings from 32 participants who viewed 40 one-minute audiovisual stimuli and provided retrospective ratings of valence and arousal after each trial [2]. We used the official preprocessed DEAP signals sampled at 128 Hz. The 60-s stimulation interval was divided into 4-s windows with a 1-s stride, resulting in 57 chronologically ordered segments per trial. The 3-s pre-stimulus reference was excluded from the segment sequence but retained for construction of the rest-relative EEG input. All 32 EEG channels and the eight DEAP peripheral channels—HEOG, VEOG, ZEMG, TEMG, GSR, respiration, plethysmography, and temperature—were retained for the subsequent EEG and PPS branches.

DREAMER was used as a second target dataset to evaluate whether the same framework extends beyond DEAP. DREAMER contains recordings from 23 participants exposed to 18 audiovisual stimuli, with EEG and ECG acquired during emotion elicitation and retrospective valence, arousal, and dominance ratings collected after each stimulus [3]. In our pipeline, the two recorded ECG channels were centered relative to the corresponding trial baseline and resampled to 128 Hz. We retained 60 s from each trial and applied the same 4-s window and 1-s stride used for DEAP, resulting in 57 temporal segments per trial. Only the two recorded ECG channels were treated as peripheral physiological inputs; unavailable DEAP-specific peripheral channels were neither zero-padded nor treated as candidate signals.

REFED plays a different role from the two target datasets. In contrast to DEAP and DREAMER, REFED provides continuous real-time valence and arousal annotations together with synchronized EEG recordings [6]. In AffectRoute, REFED supplies two external sources for TrajBridge: a REFED-supervised EEG representation used for participant-specific center adjustment and an annotation-derived temporal component used to organize within-trial variation. DEAP and DREAMER remain the target datasets for subject-independent regression under their original trial-level rating setting.

Subject independence was evaluated using participant-wise leave-one-subject-out (LOSO) validation, giving 32 outer folds for DEAP and 23 for DREAMER. In each fold, all target-dataset-dependent fitting was restricted to the remaining source participants. Throughout this study, trial coordinate refers to the prespecified trial coordinate in the acquisition protocol rather than participant identity. Because this coordinate is known before affect prediction, it is used only to retrieve the corresponding source-population trial prior. At inference, the held-out participant contributes EEG, available peripheral physiological signals (PPS), and the prespecified trial coordinate; no affect rating from that participant is used for model fitting, parameter selection, calibration, or routing. The held-out rating is accessed only after the final prediction has been frozen for evaluation.

### 2.2 AffectRoute Framework

AffectRoute divides subject-independent affect prediction into three functional sources of information instead of learning an unrestricted multimodal predictor (Fig. 1). First, a source-population trial prior provides the initial affective center for each trial coordinate using ratings from source participants only. Second, TrajBridge uses EEG together with externally supervised temporal structure derived from REFED to construct a segment-level pseudo-trajectory around this source-derived center. Third, once the TrajBridge output has been frozen, PhysioRoute evaluates whether channel-specific peripheral physiological signals (PPS) can provide a useful residual correction.

**Fig. 1.**
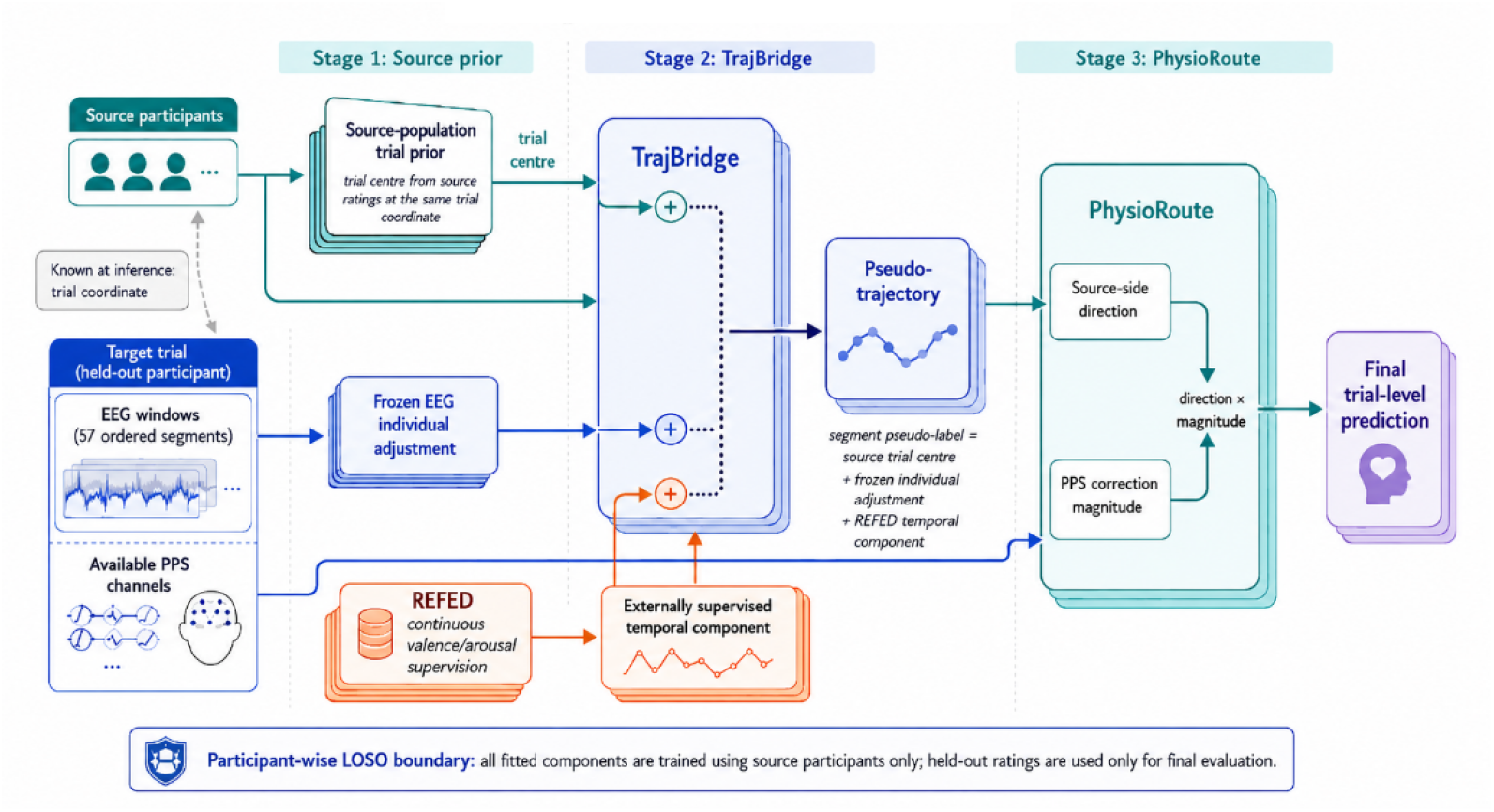
Overview of AffectRoute. A source-population trial prior establishes the initial affective center. TrajBridge uses EEG and externally supervised REFED temporal structure to construct a weakly supervised pseudo-trajectory, after which PhysioRoute evaluates source-trained channel-specific PPS residual corrections. All fitted components remain within the participant-wise LOSO boundary.

The three stages therefore have deliberately distinct roles. The trial prior represents population-level information associated with the predefined protocol, EEG provides participant-specific adjustment and supports pseudo-trajectory construction, and PPS is restricted to residual evidence rather than acting as a second complete affect predictor. All fitted components are reconstructed within each outer LOSO fold using source participants only. The framework is evaluated stage-wise as trial prior, trial prior + TrajBridge, and trial prior + TrajBridge + PhysioRoute.

### 2.3 TrajBridge: Weakly Supervised Pseudo-Trajectory Construction

Although DEAP and DREAMER provide only one affect rating for each trial, the within-trial emotional sequence is not directly observed [2,3]. Assigning the same rating to every EEG window does not represent possible within-trial variation, whereas estimating an unconstrained segment-level sequence from a single trial rating is inherently ambiguous. Similar settings have therefore been addressed using multiple-instance and weakly supervised learning frameworks [4,5]. TrajBridge addresses this problem by separating estimation of the trial-level affective center from construction of its temporal dynamics.

TrajBridge generates a weakly supervised pseudo-trajectory by using temporal dynamics determined externally, combined with emotional data from the target dataset. The external REFED dataset provides within-trial temporal organization through its continuous valence and arousal annotations, whereas the target dataset contains the data about the level of emotions from the experiment based on the center scenario of the source group and the individual EEG. As a result, the pseudo-trajectory provides the information on emotional data and the features of participants that match the data from the target dataset while including the time variation from the data set identified externally.

#### 2.3.1 REFED-Supervised Representation and Temporal Structure

REFED provides continuous, real-time annotations of valence and arousal, enabling learning from fine-grained affective supervision [6]. TrajBridge uses REFED in two complementary ways: one path learns a REFED-supervised EEG representation from EEG signals, while the other derives a global temporal component directly from the continuous affect annotations. The two paths are constructed separately and later combined with target-domain information. They serve different purposes: the EEG representation supports participant-specific affective adjustment, whereas the annotation-derived component provides shared within-trial temporal organization.

As shown in Figure 2, the first path learns transferable EEG representations from REFED EEG recordings. This representation path uses only REFED EEG recordings. We segment the entire EEG trial into 4-second, a 1-s stride, 32-channel EEG windows. After processing the EEG windows with a common average referencing, unit conversion, and resampling, we feed the EEG windows into EEGPT to extract features [13]. We then project these features from 2048 dimensions to 200 dimensions and feed them into a supervised representation model with a timestamp-aware Transformer, a two-layer subject-set Transformer, and a MLP head.

**Fig. 2.**
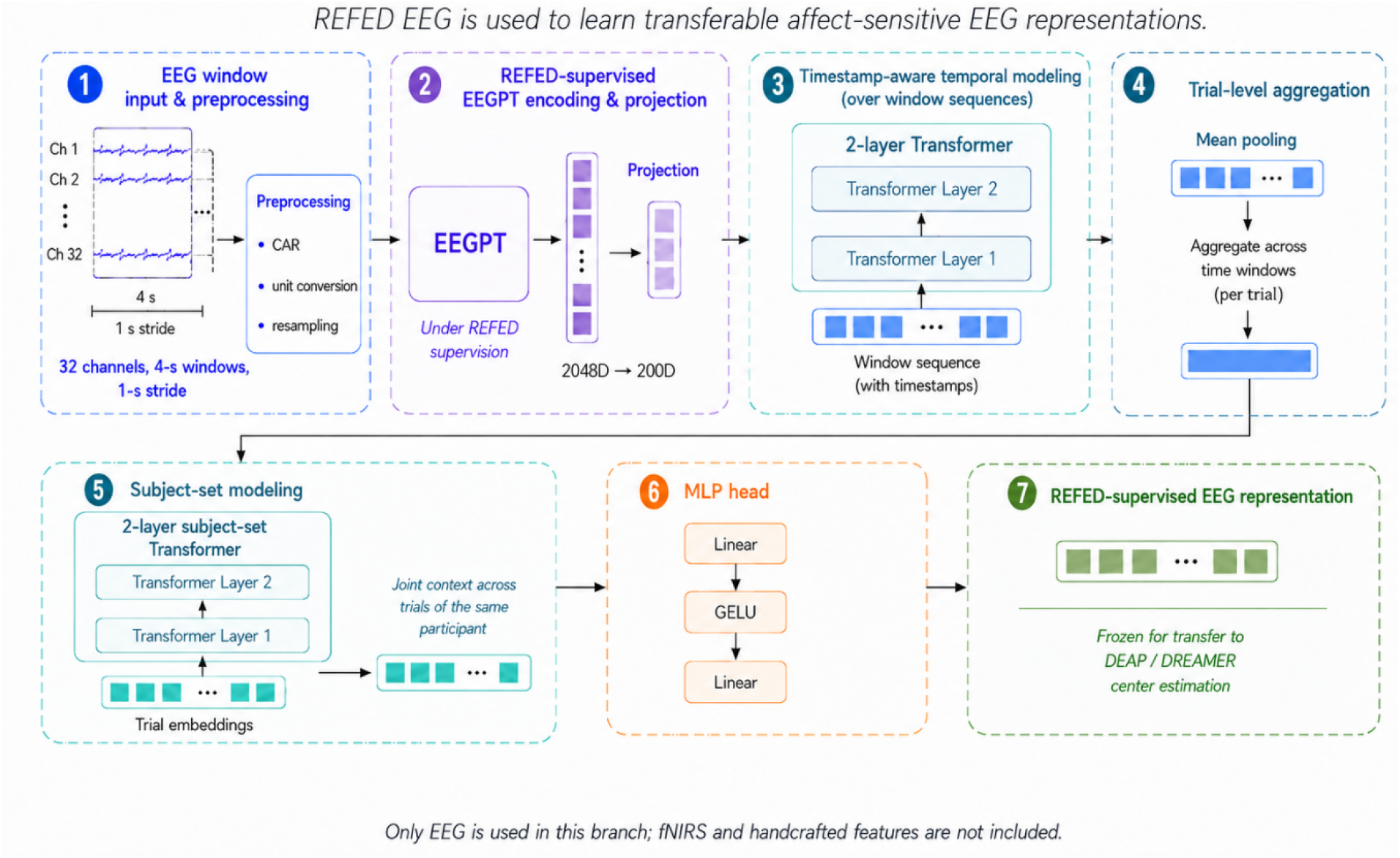
REFED-supervised EEG representation learning. REFED EEG windows are encoded using EEGPT and a supervised temporal representation model, followed by trial-level aggregation and subject-set modeling. The resulting EEG representation is frozen for transfer to DEAP and DREAMER for participant-specific center estimation.

The subject-set Transformer builds on the Set Transformer framework [14], which uses permutation-invariant attention to model structured sets. In TrajBridge, timestamps affect the estimates even though participant-specific information is not needed, because it models relationships among multiple trial representations from one person. Different from the timestamp-aware Transformer, which models the dependencies between ordered EEG windows within a trial, the subject-set Transformer operates across the trial representations from the same participant and aids in the estimation of participant-specific centers.

Developing the model with REFED involves subject-separated training, selection, and evaluation to examine transferability to the participants included in the model-fitting process. The EEG representation supervised by REFED remains fixed when features are extracted from both the DEAP and DREAMER datasets.

The second approach, illustrated in Fig. 3, derives a shared temporal component directly from continuous REFED affect annotations rather than EEG-model predictions. Previous research indicates that affective dynamics may exhibit partially shared temporal structures across individuals and naturalistic stimuli, with substantial evidence supporting the cross-movie generalizability of emotion-related representations, especially arousal [7]. For each participant and stimulus, we interpolated the data to 57 points and centered them. We then averaged the centered sequences across participants within each stimulus to generate 15 stimulus archetypes, which we subsequently averaged and re-centered by affective axis.

**Fig. 3.**
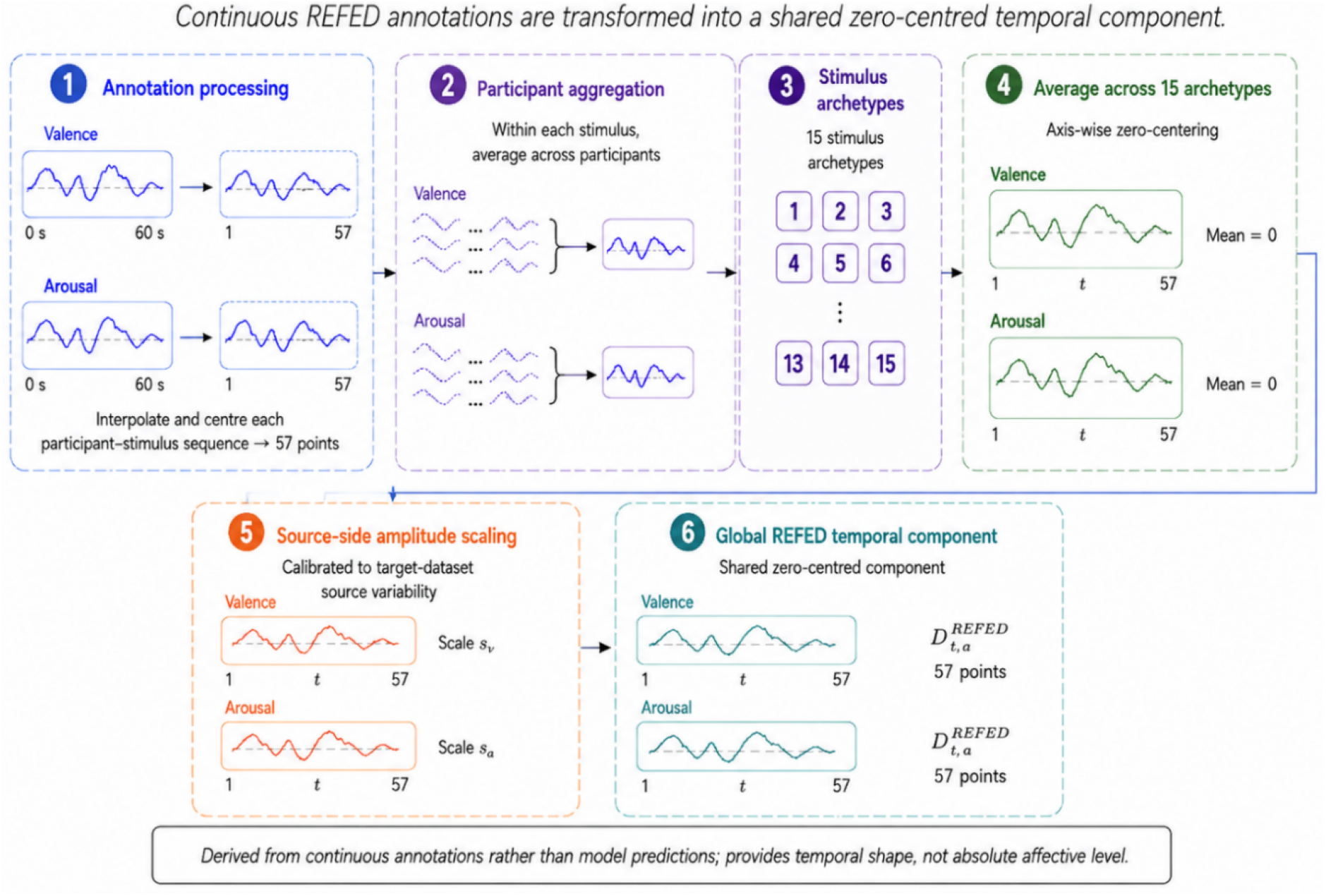
REFED-derived global temporal structure. Continuous REFED valence and arousal annotations are interpolated and centered, averaged across participants within each stimulus, and aggregated across 15 stimulus archetypes to obtain the shared zero-centered temporal component 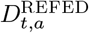. Its amplitude is subsequently calibrated using source-side information for transfer to the target datasets.

The global component obtained in this way records the relative temporal fluctuations in each trial; it serves to give the pseudo-trajectory the external supervision needed for its temporal organization, while the affective level for the target dataset is separately determined by the source population and by adjustments made to the participant on the basis of the EEG.

#### 2.3.2 Target-Domain Center Estimation and Pseudo-Trajectory Construction

Figure 4 illustrates how source-population information, transferred EEG representations, and the Global REFED temporal component are integrated to construct the TrajBridge pseudo-trajectory.

**Fig. 4.**
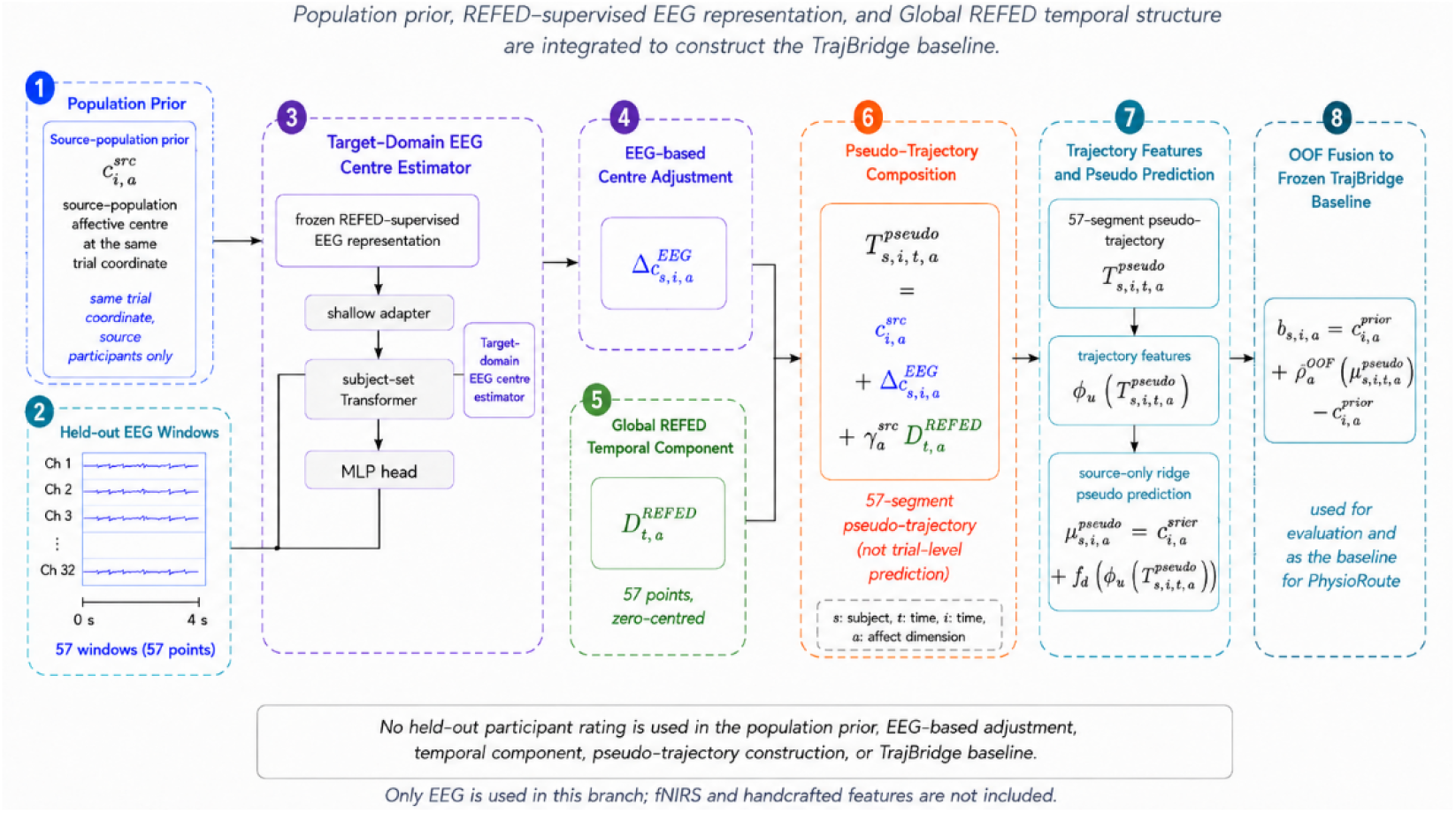
Target-domain center estimation and pseudo-trajectory construction. The source-population trial prior, the held-out participant EEG representation, and the global REFED temporal component are combined to construct the 57-segment pseudo-trajectory. The pseudo-trajectory is then summarized into trial-level trajectory features and used by a source-only ridge estimator, after which source-inner OOF fusion yields the frozen TrajBridge baseline.

For held-out participant, trial, and affective axis, a source-population center is first estimated using only ratings from source participants associated with the same trial coordinate:

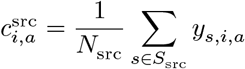

where *S*_src_ denotes the source participants within the current outer LOSO fold. The held-out participant is strictly excluded from this calculation.

The EEG recording of the held-out participant is then used to estimate the participant-specific deviation from this source-derived center. The ordered EEG windows are encoded using the frozen REFED-supervised EEG representation described in Fig. 2. A shallow target adapter adapts the transferred representation to the target dataset, and masked mean pooling aggregates window-level representations into a trial-level embedding. The trial representations from the same participant are subsequently contextualized using a subject-set Transformer, which models participant-level relationships among unlabeled trials before the MLP head estimates the individual adjustment.

The fold-specific estimator is trained exclusively on source participants and frozen at inference, producing an EEG-based adjustment 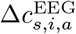, giving

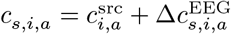

The final pseudo-trajectory combines the estimated trial center with the externally supervised temporal component derived from REFED annotations:

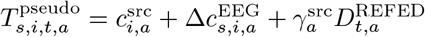

Because stimulus durations differ between REFED and the target datasets, the temporal component is transferred according to normalized within-trial position rather than absolute time. Each REFED annotation sequence is mapped from stimulus onset to offset onto a common relative time axis and interpolated to 57 temporal positions, matching the 57 target-domain segments used for DEAP and DREAMER. Here, 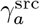is an axis-specific scaling factor estimated exclusively from source-participant EEG residuals. This source-side calibration adjusts the contribution magnitude of the REFED temporal component for the target-domain affect range while preserving the temporal organization learned from continuous affect annotations.

The 57-segment pseudo-trajectory is then converted into axis-specific trial-level trajectory features, denoted by 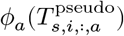. A source-only ridge estimator predicts the residual relative to the population prior 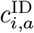 producing

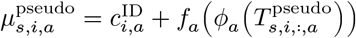

Finally, a source-inner OOF-selected fusion coefficient 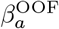 combines the pseudo prediction with the prior:

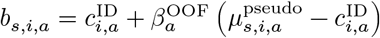

The resulting *b*_*s,i,a*_ serves as the frozen TrajBridge baseline for the subsequent PhysioRoute stage.

### 2.4 PhysioRoute: Channel-Specific PPS Residual Correction

PhysioRoute models the residual error remaining after TrajBridge and decomposes the correction into a source-derived direction and a PPS-derived magnitude. The overall procedure is summarized in Fig. 5: source-inner OOF residuals determine the correction direction, while channel-specific PPS candidates determine the correction magnitude. For source participants, the residual is defined as

**Fig. 5.**
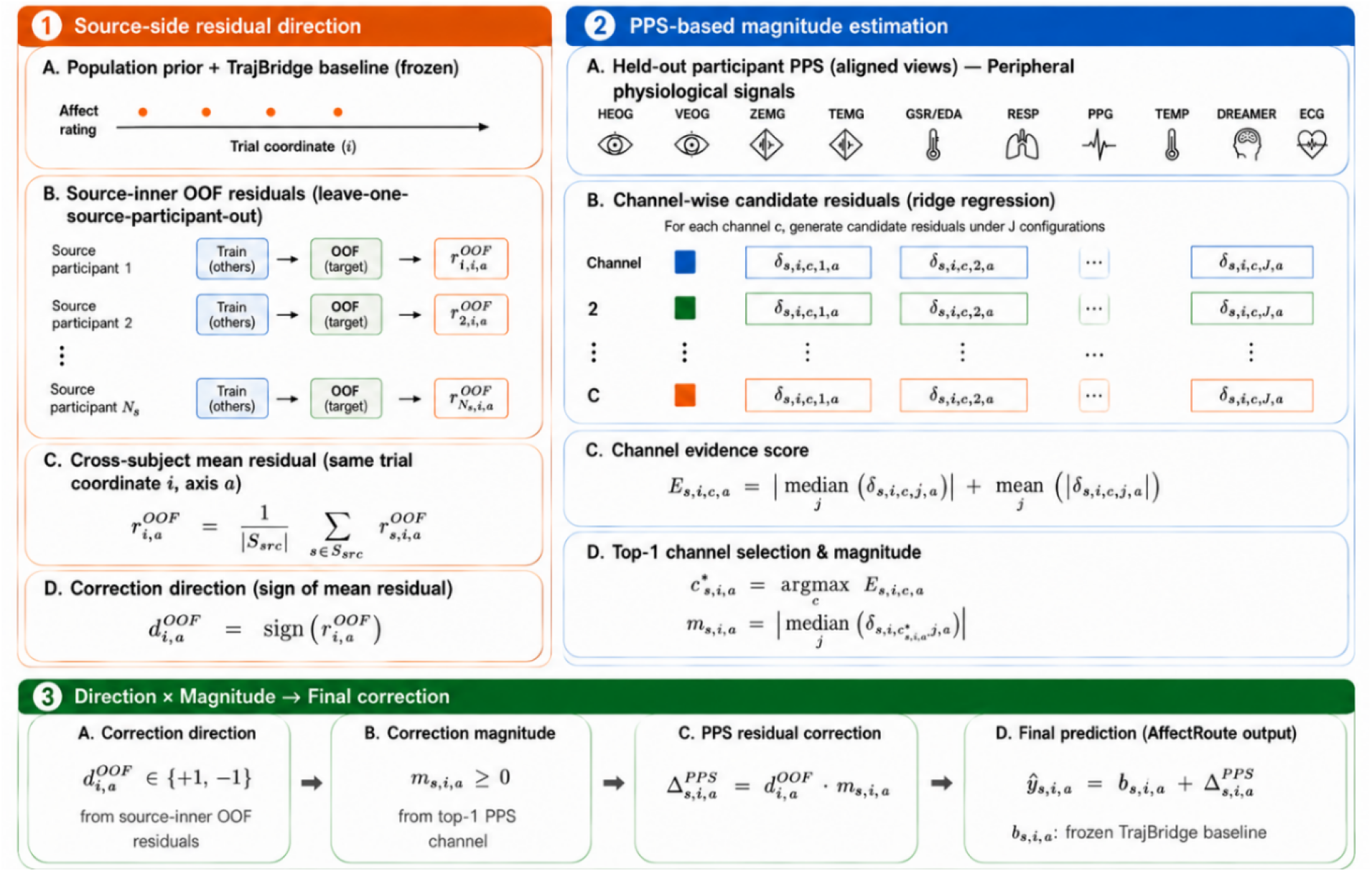
PhysioRoute channel-specific PPS residual correction. Source-inner OOF residuals determine the correction direction, while channel-specific PPS candidate residuals determine the selected correction magnitude. The resulting signed PPS residual correction is added to the frozen TrajBridge baseline to obtain the final AffectRoute prediction.

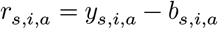

where *y*_*s,i,a*_ is the observed source-participant trial rating and *b*_*s,i,a*_ is the corresponding TrajBridge baseline prediction.

The direction of correction is determined entirely from source-inner out-of-fold (OOF) residuals. Within each outer LOSO fold, source participants are cross-fitted at the participant level so that the OOF residual of each source participant is obtained from a prediction generated without fitting on that participant where *b*_*s,i,a*_ denotes the corresponding source-inner OOF TrajBridge prediction. For a given trial coordinate and affective axis *a*, the OOF residuals from all source participants at the same trial coordinate are averaged, and the sign of this cross-subject mean defines the correction direction:

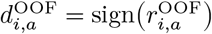

A positive value indicates that TrajBridge tends to underestimate the rating at that trial coordinate in the source fold, whereas a negative value indicates a tendency to overestimate. Thus, 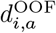 represents a same-trial cross-subject correction tendency estimated from source-inner OOF predictions.

The magnitude of the correction is estimated from channel-specific PPS evidence. Peripheral channels are represented separately rather than concatenated into a single feature block. DEAP uses all eight recorded PPS channels, whereas DREAMER uses only its two recorded ECG channels. For each channel, aligned multi-lag PPS views are combined within a single channel representation, so lag contributes as an internal feature rather than a separately selectable action. These channel representations are used to generate bounded residual candidates, and ridge regression models produce a set of candidate residual corrections:

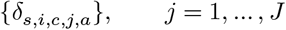

where *j* indexes a fixed set of regularization and clipping configurations. The same candidate configuration set is used throughout the LOSO evaluation and is constructed entirely on the source side, providing multiple bounded residual proposals for each channel.

PhysioRoute then performs Top-1 channel selection. For each channel, its candidate corrections are summarized by an evidence score combining their robust central tendency and overall magnitude:

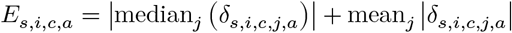

The channel with the largest evidence score is selected, and its median candidate residual determines the correction magnitude:

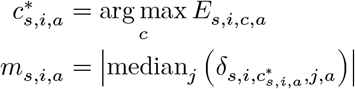

The selected magnitude is then combined with the source-derived direction to obtain the PPS residual correction and the final prediction:

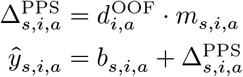

The final prediction therefore combines the TrajBridge estimate with a residual correction whose direction is derived from source-inner OOF residuals and whose magnitude is determined by the selected PPS channel.

### 2.5 Evaluation Protocol

All experiments followed the subject-independent leave-one-subject-out (LOSO) protocol described in Section 2.1. DEAP comprised 32 outer folds and DREAMER 23 outer folds. Within each fold, model fitting, parameter selection, source-inner cross-fitting, and residual construction were performed using source participants only.

Trial-level prediction performance was evaluated using mean absolute error (MAE) and concordance correlation coefficient (CCC). In addition to comparisons among the population prior, population prior + TrajBridge, and population prior + TrajBridge + PhysioRoute, paired participant-level differences were used to quantify the effect of adding PhysioRoute to the frozen TrajBridge baseline. Participant-level bootstrap resampling with 10,000 repetitions was used to estimate 95% confidence intervals.

A controlled fusion analysis compared PhysioRoute with Concatenate and Attention alternatives using the same frozen TrajBridge baseline and PPS candidate tensors.

We additionally evaluated a source-only residual control in which the same-trial source-inner OOF mean residual was added directly to the frozen TrajBridge prediction. This control allowed the source residual to determine both correction direction and magnitude without using held-out PPS evidence.

## 3. Results and discussion

This paper examines three closely related problems in subject-independent emotion regression. On the one hand, EEG- and peripheral-signal-based regression is difficult to perform in a leave-one-subject-out (LOSO) setting because emotional reactions vary widely from person to person. [1] On the other hand, both DEAP and DREAMER give only a single valence and arousal score for each trial, so the temporal sequence within each trial is not available. [2,3] Third, it may not be optimal to treat peripheral physiological signals (PPS) and EEG as interchangeable predictors when they differ in cross-subject reliability and functional role. [8-12] To address these issues, we developed AffectRoute for subject-independent affect regression. A source-only population prior serves as an initial reference point; TrajBridge combines a participant-specific EEG center adjustment with externally supervised REFED temporal structure [6] in order to create a weakly supervised segment-resolved pseudo-trajectory; and PhysioRoute breaks down the residual correction into a source-derived direction and a channel-specific PPS magnitude. Overall, these components assign separate roles to population information, EEG, and peripheral physiology rather than forcing them into a single unrestricted predictor.

### 3.1 Overall predictive gains

We first compared the source-population trial prior with a dataset-level global-mean baseline. As shown in Table 1, conditioning the population estimate on the prespecified trial coordinate reduced MAE for both affective axes in DEAP and DREAMER. The reduction was particularly pronounced for valence, decreasing MAE by 0.5766 in DEAP and 0.8601 in DREAMER, while corresponding reductions of 0.1472 and 0.4103 were observed for arousal. These results indicate that the source-population trial prior captures substantial protocol-level affective regularity and therefore provides a strong population anchor for the subsequent physiological stages.

**Table 1.**
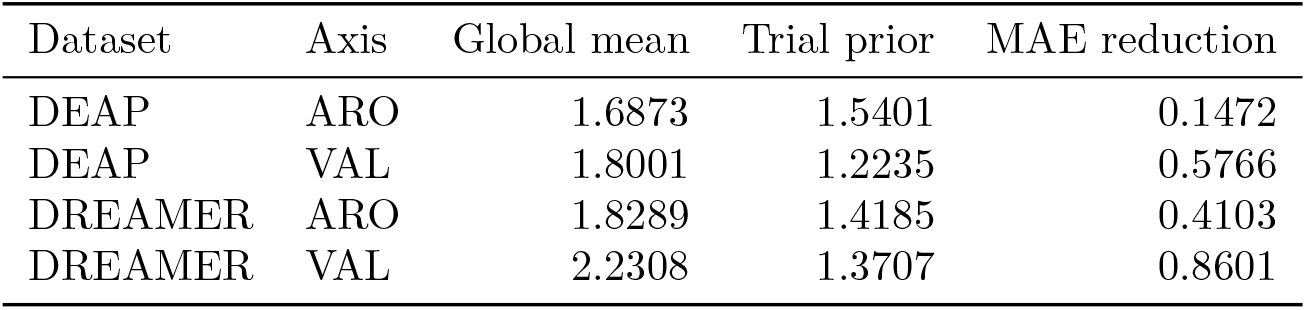
Baseline comparison between the global mean and the source-population trial prior under subject-independent evaluation.

Table 2 further decomposes the incremental contribution of TrajBridge and PhysioRoute beyond this source-population trial prior. In DEAP arousal, TrajBridge reduced MAE from 1.5401 to 1.3210 and increased CCC from 0.2795 to 0.5187, after which PhysioRoute further reduced MAE to 1.0790 and increased CCC to 0.6470. For DEAP valence, the corresponding progression was 1.2235/0.6455 for the trial prior, 1.1987/0.6595 after TrajBridge, and 0.9863/0.7427 after PhysioRoute. The same stage-wise pattern was observed in DREAMER: arousal improved from 1.4185/0.4744 to 1.3057/0.5010 and then to 1.2226/0.5757, while valence improved from 1.3707/0.7326 to 1.3071/0.7571 and finally to 1.1349/0.7979. Together, Tables 1 and 2 separate the contribution of protocol-level population information from the additional EEG-based participant information and externally supervised temporal structure introduced by TrajBridge and the subsequent PPS-based residual correction of PhysioRoute.

**Table 2.** Stage-wise performance of AffectRoute on DEAP and DREAMER under participant-wise LOSO evaluation.

| Dataset | Axis | Source-population trial prior |  | TrajBridge |  | TrajBridge + PhysioRoute |  |
| --- | --- | --- | --- | --- | --- | --- | --- |
|  |  | MAE | CCC | MAE | CCC | MAE | CCC |
| DEAP | ARO | 1.5401 | 0.2795 | 1.3210 | 0.5187 | 1.0790 | 0.6470 |
| DEAP | VAL | 1.2235 | 0.6455 | 1.1987 | 0.6595 | 0.9863 | 0.7427 |
| DREAMER | ARO | 1.4185 | 0.4744 | 1.3057 | 0.5010 | 1.2226 | 0.5757 |
| DREAMER | VAL | 1.3707 | 0.7326 | 1.3071 | 0.7571 | 1.1349 | 0.7979 |

### 3.2 Functional role separation is the central design principle

Multimodal methods for recognizing emotional states usually combine EEG with peripheral physiological signals (PPS) in a joint predictor. Cross-subject studies have examined data-, feature-, and decision-level fusion of EEG and peripheral signals [8], whereas more recent designs use modality-specific encoders with attention-based fusion to learn shared affective representations [9]. Although these approaches show the complementary nature of EEG and PPS, they still require the fusion model to decide how the heterogeneous modalities should jointly contribute to the overall affect prediction. This is difficult because physiological modalities differ in their generation mechanisms, temporal dynamics, and cross-subject reliability. Furthermore, the quality of the peripheral measurements can depend on the conditions under which they are obtained: electrodermal and photoplethysmographic signals are affected by factors such as sensor position and motion artifacts, all of which can change the quality of the signals and the reliability of the derived physiological measures [10]–[12]. As a result, considering all the available modalities as equally reliable contributors to the joint prediction might cause the final affect estimate to be influenced by unreliable or context-dependent peripheral evidence.

AffectRoute deals with this issue by giving EEG and PPS non-interchangeable predictive roles; the source-population prior serves as the starting point, TrajBridge estimates a participant-specific pseudo-trajectory by using the held-out participant’s EEG along with the externally supervised REFED temporal structure [6], and PhysioRoute then applies corrections in the direction given by the same-trial source-inner OOF residuals and using the channel-specific PPS candidates to determine the size of the corrections.

Under subject-independent evaluation, this separation of roles has two practical benefits. On the one hand, it reduces cross-modal entanglement by not requiring the heterogeneous EEG and PPS representations to be jointly mapped onto the full target; on the other hand, the direction-magnitude decomposition separates out a transferable cross-subject residual tendency from the physiological level of the correction. The methodological contribution thus consists in structured residual inference rather than in the individual PPS summary statistics.

### 3.3 TrajBridge transfers externally supervised temporal structure to weakly labeled trials

Affective responses evolve continuously over time [17], whereas DEAP and DREAMER provide only a single trial-level valence and arousal rating for each trial, leaving the within-trial affective sequence unobserved [2], [3]. It is now common practice to assign the same trial rating to every temporal segment; however, it may suppress possible within-trial variation, whereas estimating segment-level affect directly from a single post-stimulus label is inherently weakly supervised and temporally ambiguous because many different segment sequences may be compatible with the same trial-level rating [4], [5]. TrajBridge addresses this ambiguity by treating each trial as a temporally ordered sequence of EEG windows and constraining the latent segment estimates through trial-level supervision together with externally learned temporal structure.

TrajBridge creates detailed fine-grained pseudo-trajectories by combining two types of REFED transfer with information from the source in the target domain. Previous studies have shown that affective dynamics can include some common temporal features among different individuals and that certain affective representations, especially those related to arousal, can be applied across participants and naturalistic movie stimuli [15], [16], [7]. REFED provides both an EEG representation that is supervised by REFED and a temporal component which is obtained directly from the continuous valence and arousal annotations [6]. The EEG representation enables the estimation of the center specific to each participant, while the component derived from the annotations offers externally supervised temporal organization within each trial.

In each target-dataset LOSO fold, the REFED temporal component is combined with a separately estimated trial center. Source-subject ratings from the same trial define the population-level affective center, while the held-out participant’s EEG provides an individual adjustment to that center. The resulting 57-segment pseudo-trajectory therefore combines three information sources:

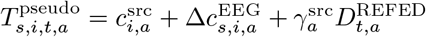

The 57-segment pseudo-trajectory is then summarized into trial-level trajectory features 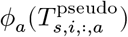, from which a source-only ridge estimator produces 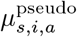. Source-inner OOF fusion then yields the frozen TrajBridge baseline 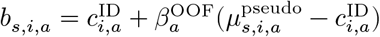, which is passed to PhysioRoute.

This construction separates where the trial is centered from how within-trial variation is organized. The trial center is determined from source-subject labels and the held-out participant’s EEG adjustment, whereas segment-wise variation is supplied by temporal structure learned independently from continuously annotated REFED data. TrajBridge therefore combines temporal context within trials with participant-level context across trials, while the population prior provides a complementary cross-participant reference for the same trial. Temporal transfer is based on relative within-trial progression rather than absolute stimulus time. In this way, target EEG supports participant-specific center adjustment and REFED provides externally supervised temporal organization, replacing uniform segment labels with a structured pseudo-trajectory.

### 3.4 PhysioRoute uses peripheral physiology for structured residual refinement

In order to determine whether the improvements achieved by PhysioRoute could be attributed entirely to the transferable residual structure from the source side, we created a control condition that consisted of adding the mean residual from the source inner OOF of the same trial directly to the frozen TrajBridge prediction, thereby letting the source residual decide both the direction and the amount of the correction. This control still proved to be clearly worse than PhysioRoute: for arousal, PhysioRoute lowered the MAE by 0.2847 and raised the CCC by 0.1554, and for valence it decreased the MAE by 0.2509 and increased the CCC by 0.1026. The results show that the benefits obtained with PhysioRoute cannot be accounted for by the residual regularity derived from the source alone and that evidence from participant-specific PPS contributes additional information to the refinement of the residual.

It is unlikely that peripheral physiological signals will contribute in a uniform way to the prediction of affect since different channels relate to separate autonomic and somatic processes and their importance may differ depending on the affective dimension and from person to person. Earlier research has found physiological associations that vary by dimension, such as the link between facial EMG and valence, between electrodermal activity and arousal, and also the cardiovascular changes that occur during emotional stimulation [17], [18]. Instead of asking the peripheral physiological signals to predict the full affective target, PhysioRoute uses the peripheral signals following TrajBridge as channel-specific information for refining the residual magnitude, the same-trial source-inner OOF residuals providing the correction direction.

The results at the channel level, as shown in Table 3, illustrate the way this physiological signal is distributed among the different sensors. The channel indices are in the same order as those used in DEAP [2]. With regard to arousal, CH04 (GSR) was chosen most often, representing 22.8% of the trials, while CH05 (RESP) yielded the highest mean MAE improvement of +0.247 even though it was only selected in 12.8% of the trials. As for valence, the selections were mainly made among CH02 (ZEMG), CH01 (VEOG) and CH06 (PPG), which were chosen in 22.7%, 21.1% and 17.6% of the trials respectively. Hence, although selection frequency and correction contribution are related, they are not the same, and the preferred physiological evidence varies between valence and arousal.

**Table 3.** DEAP PPS channel selection and contribution under strict LOSO residual correction with the TrajBridge baseline.

| Channel | Selected n (%) | Subjects (n) | MAE gain [95% CI] | Pooled CCC delta | Positive gain (%) | Direction accuracy (%) | Mean PPS magnitude |
| --- | --- | --- | --- | --- | --- | --- | --- |
| **Panel A. ARO channel selection and contribution** |  |  |  |  |  |  |  |
| CH00 | 94 (7.3) | 25 | +0.207 [+0.151, +0.258] | -0.168 | 93.6 | 93.6 | 0.405 |
| CH01 | 158 (12.3) | 30 | +0.181 [+0.122, +0.232] | -0.086 | 81.6 | 86.7 | 0.298 |
| CH02 | 171 (13.4) | 30 | +0.216 [+0.173, +0.254] | -0.109 | 88.9 | 94.7 | 0.298 |
| CH03 | 205 (16.0) | 29 | +0.216 [+0.170, +0.256] | -0.136 | 85.4 | 92.2 | 0.359 |
| CH04 | 292 (22.8) | 31 | +0.240 [+0.208, +0.274] | -0.123 | 86.3 | 91.8 | 0.325 |
| CH05 | 164 (12.8) | 27 | +0.247 [+0.222, +0.273] | -0.137 | 90.9 | 95.1 | 0.343 |
| CH06 | 120 (9.4) | 28 | +0.237 [+0.207, +0.271] | -0.138 | 90.0 | 91.7 | 0.319 |
| CH07 | 76 (5.9) | 18 | +0.214 [+0.164, +0.260] | -0.122 | 81.6 | 88.2 | 0.356 |
| TOTAL | 1,280 (100.0) | 32 | +0.242 [+0.220, +0.269] | -0.127 | 87.1 | 92.0 | 0.333 |
| **Panel B. VAL channel selection and contribution** |  |  |  |  |  |  |  |
| CH00 | 89 (7.0) | 28 | +0.180 [+0.144, +0.212] | -0.086 | 87.6 | 93.3 | 0.236 |
| CH01 | 270 (21.1) | 31 | +0.215 [+0.186, +0.247] | -0.090 | 87.8 | 94.4 | 0.286 |
| CH02 | 290 (22.7) | 31 | +0.201 [+0.173, +0.231] | -0.087 | 86.6 | 92.1 | 0.297 |
| CH03 | 90 (7.0) | 24 | +0.179 [+0.131, +0.217] | -0.105 | 92.2 | 95.6 | 0.279 |
| CH04 | 164 (12.8) | 31 | +0.170 [+0.131, +0.206] | -0.086 | 88.4 | 95.1 | 0.272 |
| CH05 | 48 (3.8) | 19 | +0.109 [+0.044, +0.161] | -0.066 | 93.8 | 93.8 | 0.215 |
| CH06 | 225 (17.6) | 31 | +0.201 [+0.162, +0.235] | -0.091 | 88.4 | 93.3 | 0.287 |
| CH07 | 104 (8.1) | 18 | +0.179 [+0.116, +0.229] | -0.081 | 89.4 | 93.3 | 0.318 |
| TOTAL | 1,280 (100.0) | 32 | +0.212 [+0.199, +0.227] | -0.088 | 88.4 | 93.7 | 0.283 |

The analysis of eight channels, as shown in Fig. 6, reinforces this routing profile by showing the impact of removing each individual channel. When ZEMG, VEOG, and PPG are removed, valence is most affected, while the removal of GSR, RESP, TEMG, and HEOG has the greatest effect on arousal. the largest axis-specific effects can be understood in physiological terms, especially the role of ZEMG in valence and that of GSR in arousal. Yet the highest exclusion penalty is still relatively small, which shows that the information from the PPS is spread out among the channels and that there is a good degree of compensability rather than being concentrated in one sensor that is essential. This feature of the channels being distributed is well suited to PhysioRoute’s Top-1 routing design, since that design permits the physiological source determining the magnitude of the correction to differ from trial to trial and with respect to the affective axes.

**Fig. 6.**
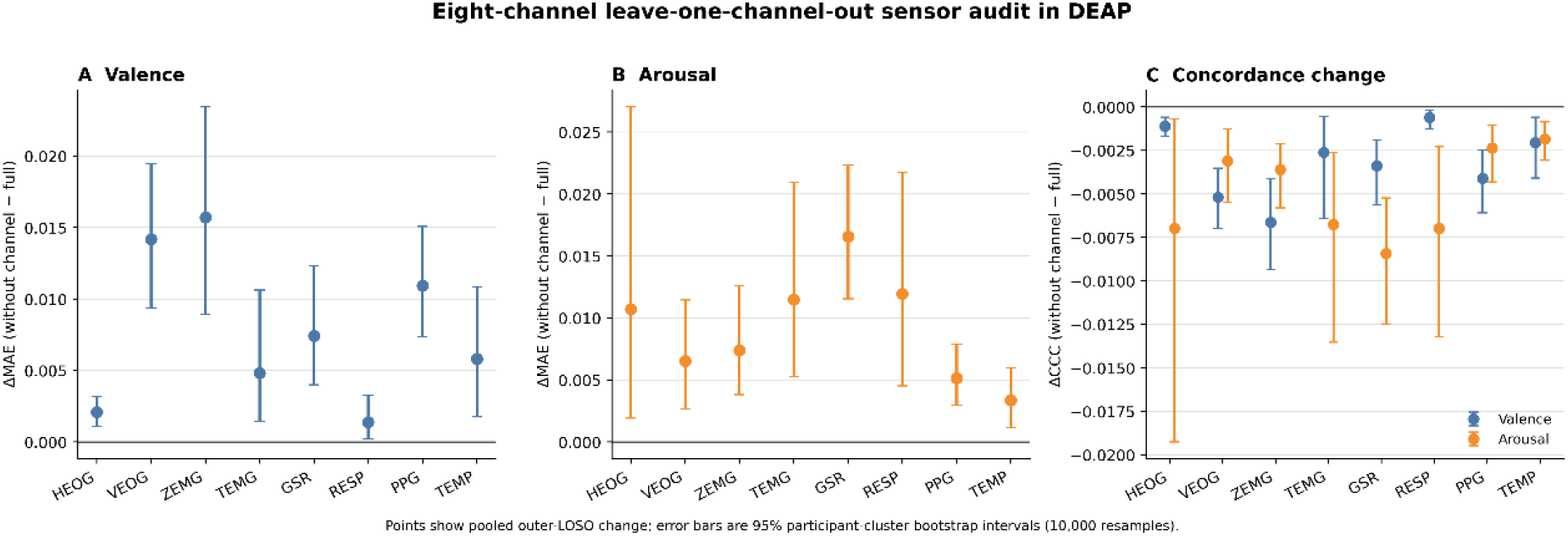
Leave-one-channel-out analysis of peripheral contributions in DEAP. A–B, Change in MAE after exclusion of each PPS channel for valence and arousal. C, Corresponding change in CCC. Positive ΔMAE and negative ΔCCC indicate performance loss after channel removal. Error bars denote 95% participant-cluster bootstrap confidence intervals based on 10,000 resamples.

The fusion ablation further clarifies the advantage of the role-structured PhysioRoute design over direct PPS fusion. Using the same TrajBridge baseline, both Concatenate and Attention were consistently inferior to PhysioRoute for all 32 held-out DEAP participants on both affective axes. For arousal, Concatenate and Attention increased MAE by 0.2556 and 0.2473, respectively, while reducing CCC by 0.1489 and 0.1469 relative to PhysioRoute. Similar differences were observed for valence, with MAE increases of 0.2261 and 0.2244 and CCC reductions of 0.0887 and 0.0899. The source-side selection behaviour further suggests that unrestricted PPS fusion was difficult to support reliably: the strongest ridge penalty (λ = 1000) was selected in nearly all source-OOF folds, effectively suppressing the direct PPS contribution, while the uniform-attention configuration (τ = 0) was selected in 24 of 32 valence folds. Together, these findings indicate that the advantage of PhysioRoute does not arise simply from adding peripheral information, but from constraining how that information enters the prediction: the frozen TrajBridge estimate is corrected using a source-derived direction and channel-specific PPS magnitude rather than allowing PPS to form an unrestricted second predictor.

### 3.5 Limitations and future directions

Even though the results are promising, there are still some limitations in this study. Firstly, both DEAP and DREAMER provide only retrospective trial-level ratings, not independent, segment-level continuous annotations. [2,3] It is therefore only possible for TrajBridge to create a weakly supervised pseudo-trajectory, its effectiveness being mainly verified by the trial-level predictions and by externally provided temporal structures. Because the target datasets do not include true segment-level labels, we cannot directly assess whether the underlying moment-to-moment affective trajectory has been accurately captured.

Second, our evaluation makes use of only two commonly used, laboratory-based affective datasets. [2,3] Although DEAP and DREAMER do provide complementary EEG and peripheral physiological recordings, both of these studies involve participant groups that are relatively homogeneous and elicit emotions in a laboratory setting. If we are to properly assess the generalizability of our framework, further research will have to include larger and more diverse groups as well as more naturalistic recording environments.

Thirdly, the present evaluation concentrates on new participants who are within the predefined acquisition situation in which the trial coordinate is available. It is still an important area for future research to extend AffectRoute to unseen stimuli and to carry out affect estimation in open contexts.

## 4. Conclusion

AffectRoute separates two tasks that are often combined in subject-independent multimodal affect regression: constructing a fine-grained EEG-centered affect estimate from coarse trial-level supervision, and using peripheral physiology to refine the remaining residual. TrajBridge combines a source-population anchor, participant-specific EEG adjustment, and REFED-derived temporal organization to form a weakly supervised segment-resolved pseudo-trajectory. PhysioRoute then uses a source-derived correction direction together with channel-specific PPS magnitude to refine the TrajBridge estimate.

Across DEAP and DREAMER, the source-population trial prior improved on a global-mean baseline, and both TrajBridge and PhysioRoute provided further stage-wise gains. Channel-level selection and leave-one-channel-out analyses showed that PPS contributions are axis-dependent and distributed across multiple sensors, while fusion comparisons showed that structured peripheral residual refinement provides a consistent alternative to unrestricted multimodal fusion. These results support assigning population information, REFED annotation-derived / externally supervised temporal structure and participant context, and peripheral physiology distinct roles within the prediction pipeline. Future work can extend the pseudo-trajectory analysis to target datasets with continuous annotations and broader stimulus settings.

